# Realtime 2-5A kinetics suggests interferons β and λ evade global arrest of translation by RNase L

**DOI:** 10.1101/476341

**Authors:** Alisha Chitrakar, Sneha Rath, Jesse Donovan, Kaitlin Demarest, Yize Li, Raghavendra Rao Sridhar, Susan R. Weiss, Sergei V. Kotenko, Ned S. Wingreen, Alexei Korennykh

## Abstract

Cells of all mammals recognize double-stranded RNA (dsRNA) as a foreign material. In response, they release interferons (IFNs) and activate a ubiquitously expressed pseudokinase/endoribonuclease RNase L. RNase L executes regulated RNA decay and halts global translation. Here we developed a biosensor for 2’,5’-oligoadenylate (2-5A), the natural activator of RNase L. We found that 2-5A was acutely synthesized by cells in response to dsRNA sensing, which immediately triggered cellular RNA cleavage by RNase L and arrested host protein synthesis. However, translation-arrested cells still transcribed IFN-stimulated genes (ISGs) and secreted IFNs of types I and III (IFN-β and IFN-λ). Our data suggests that IFNs escape from the action of RNase L on translation. We propose that 2-5A/RNase L pathway serves to rapidly and accurately suppress basal protein synthesis, preserving privileged production of defense proteins of the innate immune system.

**Significance:** RNase L is a mammalian enzyme that can stop global protein synthesis during interferon response. Cells must balance the need to make interferons (which are proteins) with the risk to lose cell-wide translation due to RNase L. This balance can most simply be achieved if RNase L was activated late in the interferon response. However, we show by engineering a biosensor for the RNase L pathway, that on the contrary, RNase L activation precedes interferon synthesis. Further, translation of interferons evades the action of RNase L. Our data suggest that RNase L facilitates a switch of protein synthesis from homeostasis to specific needs of innate immune signaling.

## Introduction

Interferons IFNs of type I (α and β) and type III (λ1, λ2 and λ3) are cytokines secreted by cells after exposure to pathogens or internal damage. Both types of IFNs activate strong transcriptional programs in surrounding tissues and have a central role in the innate immune system. Production of the IFNs requires protein synthesis. However, during innate immune response to double-stranded RNA (dsRNA), IFN production is universally accompanied by translational arrest. Inhibition of protein synthesis arises in part due to activation of the dsRNA-dependent protein kinase R (PKR), and in part due to signaling by conserved small RNAs that contain 2’,5’-linked oligoadenylates (2-5A)^1-4^ The action of 2-5A is sufficient for arrest of translation, independent of PKR, and at least in some cell lines 2-5A is the main cause of translational arrest^5^. In human cells, 2-5A is synthesized by three enzymes: oligoadenylate synthetase 1 (OAS1), OAS2 and OAS3 (OASs), which function as cytosolic dsRNA sensors using dsRNA binding for activation^6,7^. The activity of the OASs is normally low to allow housekeeping protein synthesis, but it increases in the presence of viral or host dsRNA molecules^8^. 2-5A has a strong antiviral effect, against which many viruses have evolved 2-5A antagonist genes that are essential for infection^9-16^.

The 2-5A system is also a surveillance pathway for endogenous dsRNAs from mammalian genomes. Cells with adenosine deaminase 1 (ADAR1) deficiency accumulate self-dsRNA that promotes 2-5A-driven apoptosis^8^. 2-5A synthesis in the presence of low amounts of endogenous dsRNAs does not cause cell death, but functions as a suppressor of adhesion, proliferation, migration, and prostate cancer metastasis^17,18^. Additionally, activation of the 2-5A system also blocks secretion of milk proteins and stops lactation, presumably as a mechanism to prevent passing infection via breast feeding^19^.

All of the effects of 2-5A arise from the action of a single mammalian 2-5A receptor, pseudokinase-endoribonuclease L (RNase L)^20^. 2-5A binds to the ankyrin-repeat (ANK) domain of RNase L and promotes its oligomerization and formation of a dimeric endoribonuclease active site^21-23^. This dimer further assembles into high-order oligomers^24^ that cleave viral RNAs^16,18^ and all components of the translation apparatus, including mRNAs^25^, tRNAs^26^ and 28S/18S rRNAs^27,28^. The resulting action of RNase L inhibits global translation, which puts all proteins, including IFNs, at risk of arrest during a cellular response to dsRNA. The impact of translational shutdown by RNase L on IFN synthesis and paracrine IFN signaling is unknown. Measurements of 2-5A/RNase L activity have been limited by the need for biochemical analysis, which are incompatible with live cells. To address this challenge, we developed a realtime 2-5A biosensor and used it to elucidate the kinetics of 2-5A-mediated RNase L activation and translational arrest taking place during the cellular response to immuno-stimulatory dsRNAs. Our biosensor can detect in situ 2-5A synthesis in mammalian cells and thereby it establishes a heretofore missing platform for cell-based applications. These applications can range from live cell screens for modulators of innate immune responses to mechanistic analysis of dsRNA sensing, which we describe in our present work.

## Results

### Realtime 2-5A dynamics in live cells

In the absence of methods to monitor 2-5A without cell disruption, the cellular dynamics of this second messenger are poorly understood. Here we developed a biosensor for continuous and non-invasive 2-5A monitoring in live cells. The biosensor was designed based on the crystal structures of the cellular 2-5A receptor RNase L^22,24^ (Fig. 1A), which indicate that the N-terminal ANK domain of RNase L is sufficient for 2-5A sensing and provides a minimal dimerization module^24^. In a single ANK protomer, the N- and C-termini in *cis* are separated, but upon dimerization of two ANK domains, which bind notably head-to-tail, the C-/N-termini become positioned in *trans* next to each other (Fig. 1B). We employed this in *cis* vs *in trans* N-/C-distance decrease to drive a dual-split-luciferase sensor.

**Fig. 1.**
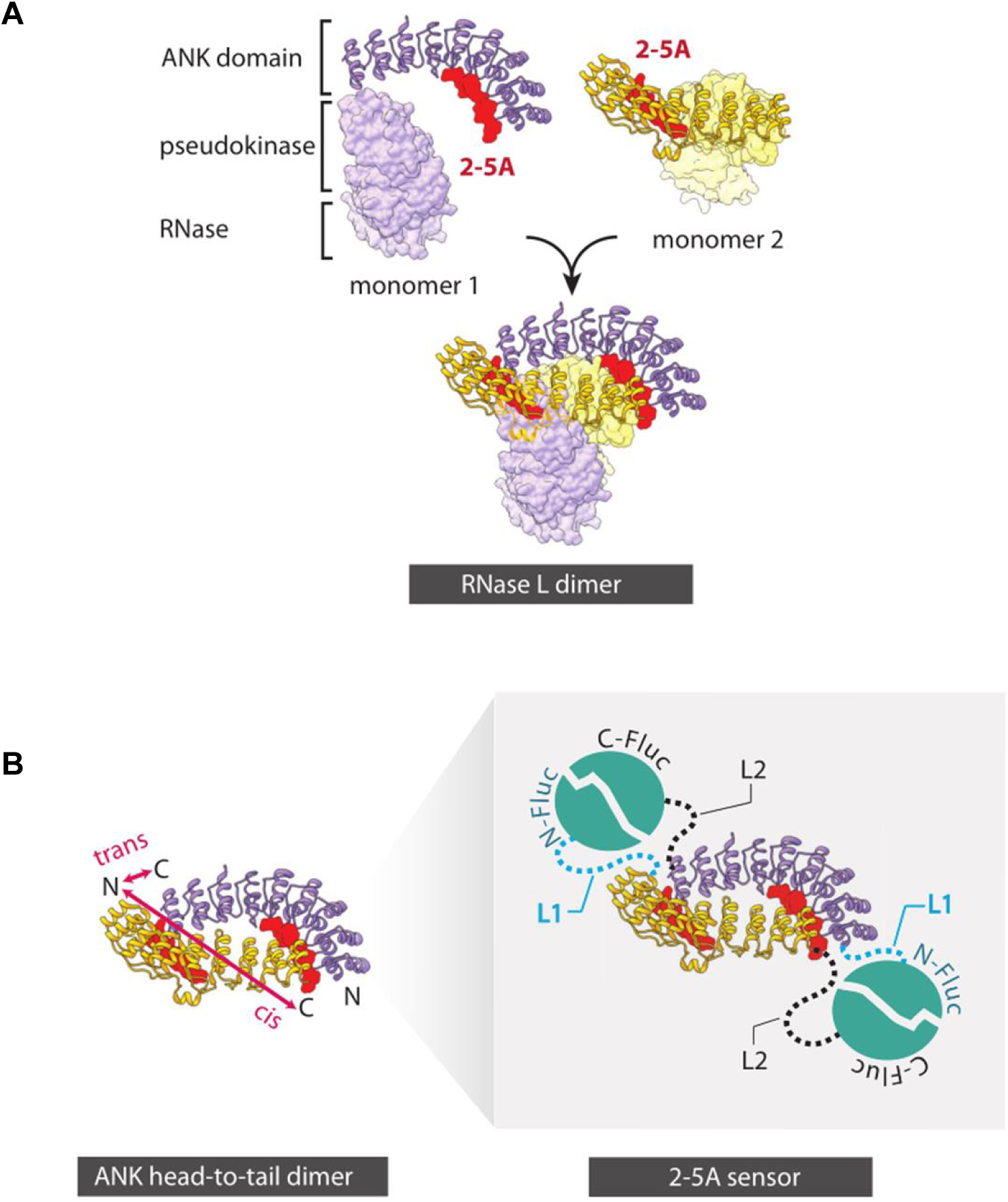
Structure-guided design of the 2-5A biosensor. (**A**) The N-terminal ANK domain of RNase L can bind two molecules of 2-5A (red) and form a 2-5A-templated homodimer with a head-to-tail structure. (**B**) In the 2-5A biosensor, split firefly luciferase halves are engineered on both termini of the ANK domain. Upon 2-5A binding, the head-to-tail configuration of the ANK domains brings the N- and C-termini of two different ANK domains in proximity, which reconstitutes two functional luciferase copies per assembled ANK/ANK reporter. Activation of luminescence provides the readout of 2-5A.

The combinatorially optimized sensor has two halves, which consist of the core ANK domain fused to a modified split firefly luciferase (Fluc)^29^ (Fig. 1B). We modified Fluc by replacing the published overlapping split junction N416/C415 with a non-overlapping junction (Fig. 2A). Due to the head-to-tail structure, the sensor was encoded as one polypeptide, which simplified its use compared to usual two-protein split systems. We optimized the reporter by engineering the linker regions and selected variant V6 with the highest luminescence response to 2-5A for further work (Fig. 2A). We determined that V6 was suitable for 2-5A detection over a range of 2-5A and V6 concentrations from ~ 10 nM to at least 1 μΜ (V6) and 3 μΜ (2-5A) (Fig. 2B-C). This luminescence response was specific and abolished by a control mutation Y312A, which removed the key 2-5A-sensing tyrosine^24^ (Fig. 2B-C). The reporter V6 had sufficient sensitivity to detect low amounts of 2-5A purified from human cells treated with poly Inosine/poly-Cytidine (poly-IC) dsRNA (Fig. S1A). These reporter measurements closely agreed with the standard endoribonuclease readout based on RNA cleavage (Fig. S1B).

**Fig. 2.**
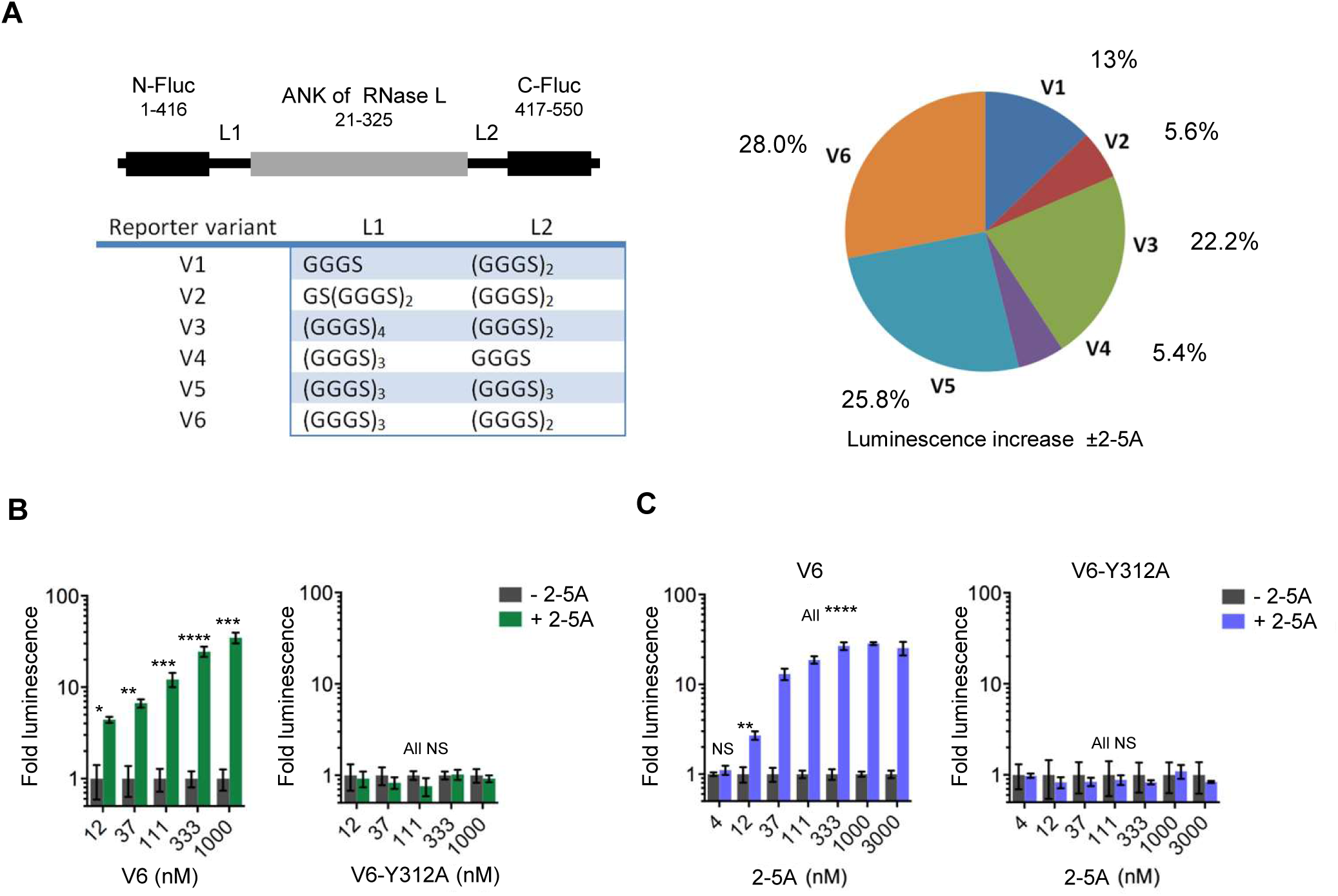
Light-based detection of nanomolar 2-5A concentrations. (**A**) Luminescence analysis of 2-5A biosensor variants (left) in A549 cells. Data on their relative activation by synthetic 2-5A (right) are from three independent experiments (two for V1, V3 and V4). (**B** and **C**) Luminescence analysis of 2-5A biosensor V6 dose response in A549 cells at excess 2-5A (B) or in response to varying concentrations of 2-5A (C). The Y312A mutant serves as a control. Data are means ± S.E. pooled from at least three independent experiments. Stars here and throughout text show statistical significance (P) from Welch two-tailed unpaired t-test (James McCaffrey implementation, Microsoft, https://msdn.microsoft.com/en-us/magazine/mt620016.aspx). *P<0.05, **P<0.01, ***P<0.001, *P≤0.0001, NS: non-significant.

To determine whether V6 was sufficiently bright and stable in live cells, we expressed FLAG-tagged V6 in HeLa cells and stimulated these reporter cells with poly-IC. Poly-IC treatment conditions were selected to produce cleavage of 28S rRNA in the conventional end-point assay using cell disruption^22^. 28S rRNA cleavage was noticeable after one hour of poly-IC treatment and further increased after three and four hours (Fig. 3A). HeLa cells expressing WT FLAG-V6 exhibited a time-dependent increase in luminescence in the presence of cell-permeable D-luciferin ethyl ester (Fig. 3B and fig. S1C). The luminescence increase began at ~ 30 minutes of poly-IC treatment and prior to appearance of visible 28S rRNA cleavage. HeLa cells expressing control FLAG-V6-Y312A produced no increase in luminescence, confirming that WT V6 detects cellular 2-5A in real time.

**Fig. 3.**
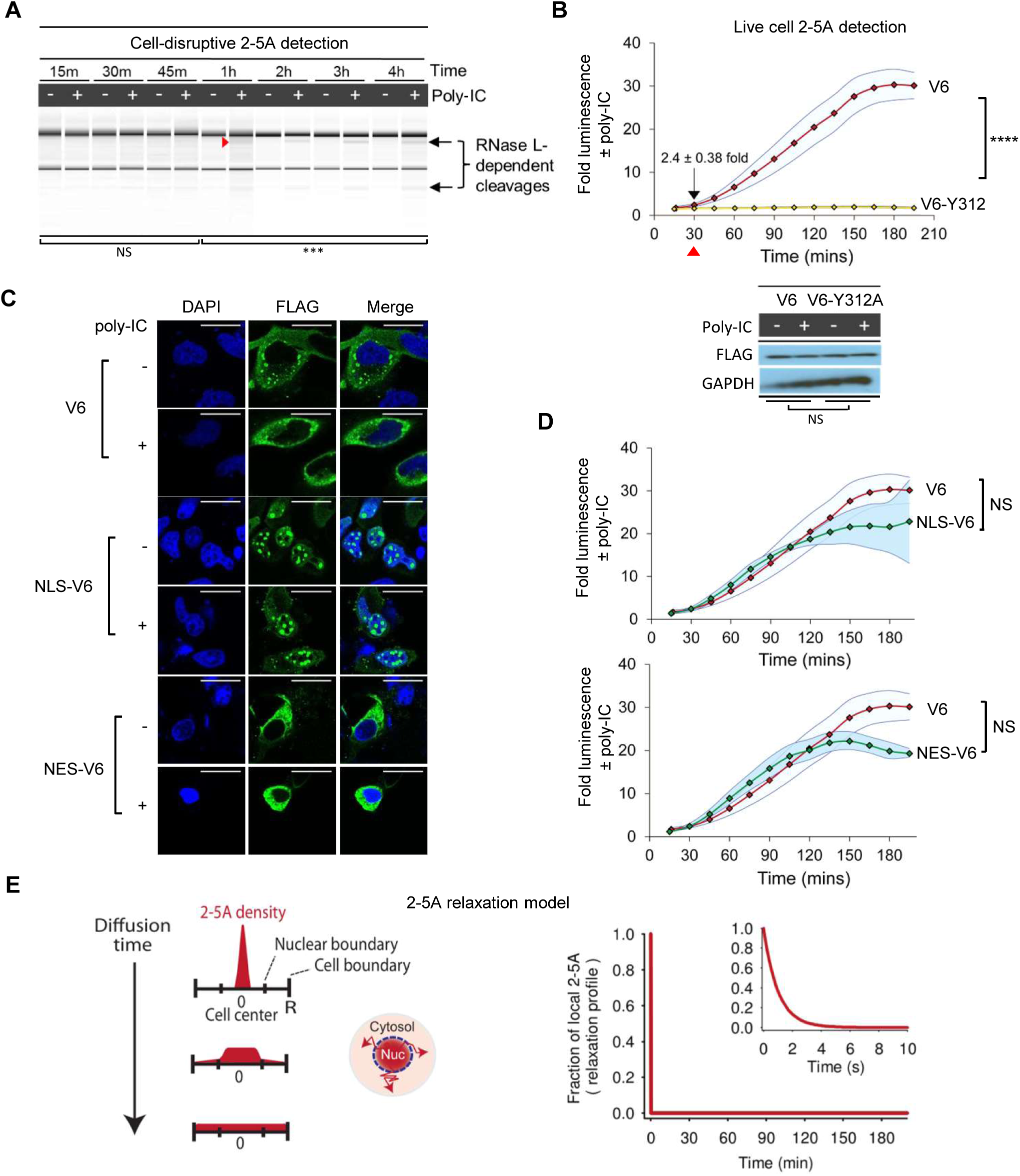
Continuous 2-5A monitoring in live cells. (**A**) RNA nano-chip analysis of total RNA degradation in HeLa cells treated with poly-IC for the indicated times. Images are representative of 3 independent experiments. Statistical significance is shown for aggregate “+” vs “-“ lines. (**B**) Realtime luminescence analysis of 2-5A synthesis in HeLa cells transiently transfected with the V6 2-5A biosensor and poly-IC. Data (upper) are means ± S.E. pooled from three independent experiments. Immunoblot (inset) of FLAG-V6 and FLAG-V6-Y312A expression. Expression of V6 is compared to expression of Y312A as NS. Similar V6 and Y312A expression was also concluded from basal luminescence in the top panel (P-val = 0.16, NS, from 3 experimental replicates). (**C**) Immunofluorescence microscopy analysis of FLAG-V6 2-5A biosensor tagged with NLS and NES. Nuclei were stained with DAPI. Images are representative of at least two independent experiments. Scale bars, 25 μm. (**D**) Luminescence analysis of 2-5A synthesis in Hela cells expressing NLS-tagged (upper) and NES-tagged (lower) 2-5A V6 biosensor. Untagged biosensor responses are overlaid for comparison. Data are means ± S.E. pooled from 3 independent experiments. (**E**) Model of the kinetics of slowest-mode 2-5A relaxation by diffusion in cells with a nucleus, where f(t) = exp(−0.25⋅D(4.49/R)^2^⋅t). D is the 2-5A diffusion coefficient and R is the cell radius.

Further evidence for specific 2-5A detection was obtained in cells with 2-5A synthetases knocked out. It has been shown that knockout of OAS3 is sufficient to inhibit 2-5A synthesis in human cells^11^. We found that biosensor exhibited a robust response to poly-IC in OAS1-KO and OAS2-KO cells. In contrast, the response was lost in OAS3-KO cells (fig. S2), providing a confirmation that 2-5A synthesis gave rise to the reporter activity. RNase L may be activated by local production of 2-5A^30^, at secrete sites inside cells (the cytosol). Notably, OASs are present not only in the cytosol, but also in the nucleus^31-33^, which further supports that 2-5A accumulation could be non-uniform with subcellular spaces. The availability of live cell 2-5A sensor offers an opportunity to evaluate 2-5A accumulation in individual cellular compartments in situ.

Toward this end, we engineered tagged versions of V6 with nuclear localization signal (NLS) and nuclear export signal (NES). Both variants localized to the expected sites (Fig. 3C). In the presence of poly-IC, NLS-V6 and NES-V6 produced nearly identical luminescence profiles, which are similar to the trace obtained with untagged V6 (Fig. 3D). This similarity is most simply reconciled with a model of rapid 2-5A equilibration between the two compartments due to diffusion. Indeed, 2-5A produced at the center of a HeLa cell, may take only several seconds to diffuse across the nucleus, through the nuclear pores, and to become evenly distributed between the nucleus and the cytosol (Fig. 3E). Our measurements did not support localized 2-5A action and indicated that 2-5A was poised to establish communication between the OASs and RNase L across the cell.

### Translational arrest by 2-5A precedes the IFN response

The pathways of 2-5A and IFNs are closely interconnected. IFNs stimulate 2-5A production by transcriptionally inducing the OASs^3,34,35^. Conversely, 2-5A can amplify^36^ and suppress^37^ IFN-β protein production. Considering that RNase L stops translation and ultimately causes apoptosis^38^, IFNs may critically require mechanisms to delay RNase L activation and evade RNase L. To test whether such mechanisms exist, we generated stable A549 and HeLa human cell lines that carry FLAG-V6 (fig. S3) and used these cells to measure 2-5A synthesis throughout dsRNA response. Time-dependent 2-5A synthesis was readily observed over a range of poly-IC concentrations (Fig. 4A). Accumulation of 2-5A started nearly immediately after dsRNA addition, exhibiting a discernible lag at low doses and no lag at higher doses of poly-IC. In contrast, transcription of IFN stimulated genes (ISGs) measured by qPCR of OAS1/2/3/L and the helicases RIG-I and MDA5, developed with a lag of 2-4 hours and became strong after maximal 2-5A production (Fig. 4, A and B). Rapid 2-5A synthesis before the IFN response was confirmed by cleavage of 28S rRNA in A549 cells (Fig. 4C), and in HeLa cells using a combination of biosensor and qPCR readouts (fig. S4). These observations suggest that 2-5A production may precede the IFN response and that 2-5A is supplied by basal rather than IFN-induced OASs. Therefore, 2-5A/OAS activation does not require IFN stimulation, as reported^39^. Similarly, basal OASs are solely responsible for protection of mouse myeloid cells from murine coronavirus^40^. We also found that the OAS/RNase L activation was not inhibited by pre-treatment with a transcription inhibitor Actinomycin D (fig. S5) and priming the cells with IFN-β had only a modest ≤ 2-fold effect on 2-5A synthesis while a strong overall transcriptional response was present (Fig. 4, D to E, and fig. S6 and S7). These data further support the involvement of the basally expressed OASs in 2-5A production, prior to ISG transcriptional response.

**Fig. 4.**
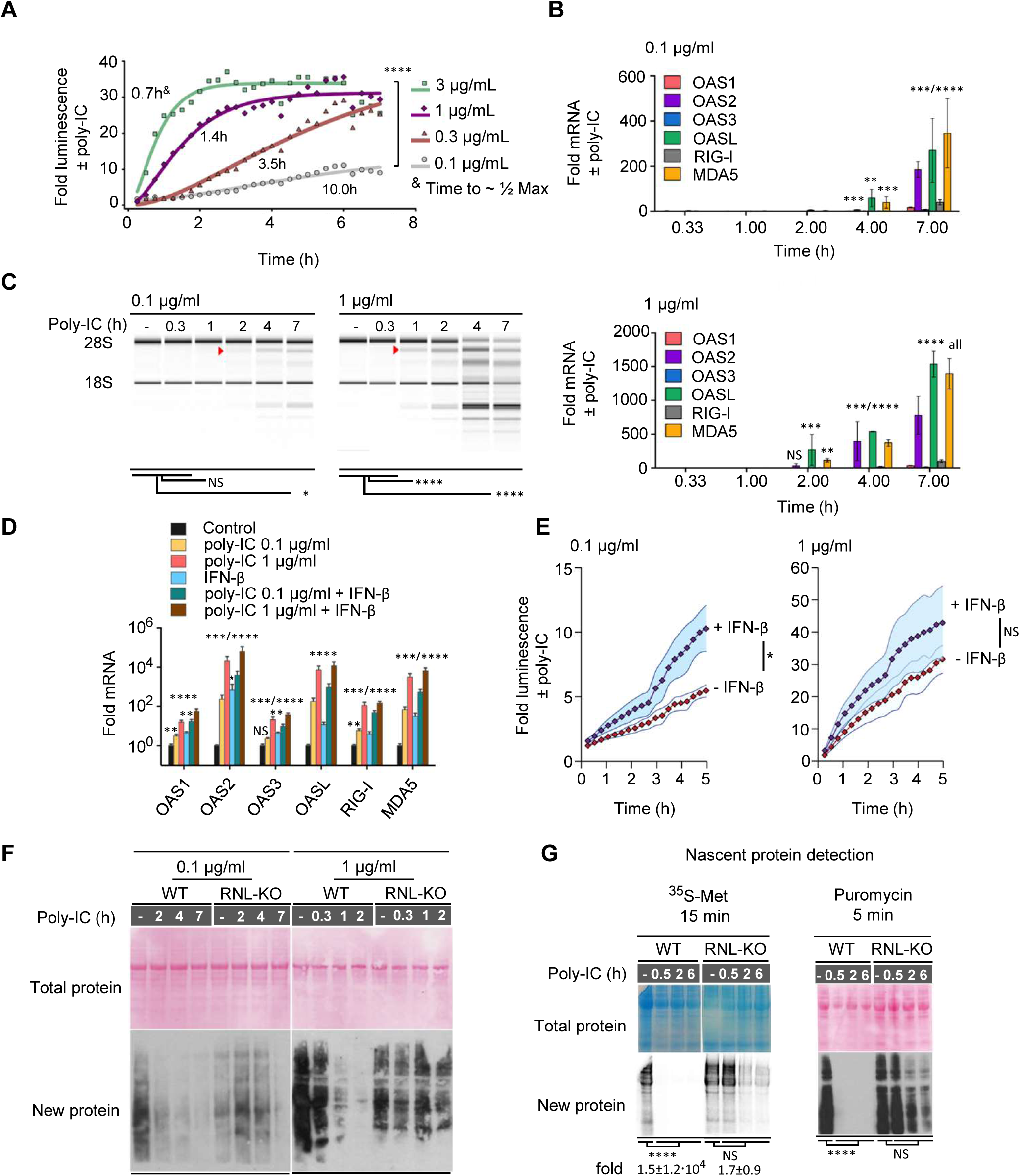
Dynamics of 2-5A, transcriptional IFN response and translation in A549 cells. (**A**) Luminescence analysis of 2-5A dynamics in A549 cells stably expressing FLAG-V6 2-5A biosensor at the indicated times after poly I:C treatment. P-value was computed from the three experiments at the highest dose of poly-IC vs 0.1 μg/ml poly-IC. (**B**) qRT-PCR analysis of ISG expression in A549 cells after poly-IC treatment vs untreated controls. Data are means ± S.E. from 3 biological replicates (4h time point with 1 μg/ml poly-IC had 2 replicates; several measurements use 4 replicates). (**C**) RNA nano-chip analysis of 28S rRNA cleavage in A549 cells treated with poly IC for the indicated times. Arrows indicate a major RNase L-induced cleavage product. Images are representative of 3 independent experiment. (**D**) qRT-PCR analysis of ISG expression in A549 cells at 24 hours after poly-IC, IFN-β, or combined treatment. Data are means ± S.E. from three biological replicates. (**E**) Luminescence analysis of 2-5A dynamics in A549 cells with and without 24-hour IFN-β pre-treatment. Data are means ± S.E. pooled from at least 3 independent experiments. (**F**) Puromycin western blot analysis of nascent protein synthesis in WT and RNase L knockout (RNL-KO) A549 cells after treatment with poly I:C for the indicated times. Blots are representative of three independent experiments. (**G**) Western blot and autoradiography analysis of nascent protein synthesis in A549 cells labeled with puromycin or ^35^S metabolic labeling after treatment with poly I:C for the indicated times. Blots are representative of four independent experiments.

To determine whether cellular 2-5A dynamics corresponds to a rapid arrest of translation by RNase L, we measured nascent protein synthesis by puromycin pulse labeling in WT and RNase L^-/-^ A549 cells^5^. Treatment of WT, but not RNase L^-/-^ cells with poly-IC halted global translation before ISG induction (Fig. 4F and 4B). RNase L^-/-^ cells exhibited a delayed and incomplete translational attenuation, presumably due to PKR. Translational arrest ahead of ISG induction was also present in HeLa cells (Fig. S4, B and C). Disengagement of basal protein synthesis before the IFN response was further confirmed using metabolic labeling of nascent proteome with ^35^S (Fig. 4G).

### IFNs β and A escape the translational shutoff caused by 2-5A

2-5A rapidly stops cell-wide protein synthesis. To examine IFN protein production under these conditions, we treated WT and RNase L^-/-^ cells with poly-IC and assayed the media for IFN activity (Fig. 5A). These tests revealed a time-dependent increase of media antiviral activity and media ability to induce ISGs, which developed after RNase L-mediated translational arrest (Fig. 5B, and fig. S8). At time points well beyond translational arrest, we observed an increase in antiviral activity (Fig. 5C) and ISG induction by two orders of magnitude from media of poly I:C-treated WT and RNase L^-/-^ cells (Fig. 5D). In agreement with the ISG induction readout, media from WT and RNase L^-/-^ cells exhibited comparable antiviral activity and similar time-dependent increase of IFN potency (Fig. S9). These data suggest that shutdown of bulk translation by RNase L thus does not inhibit IFN production.

**Fig. 5.**
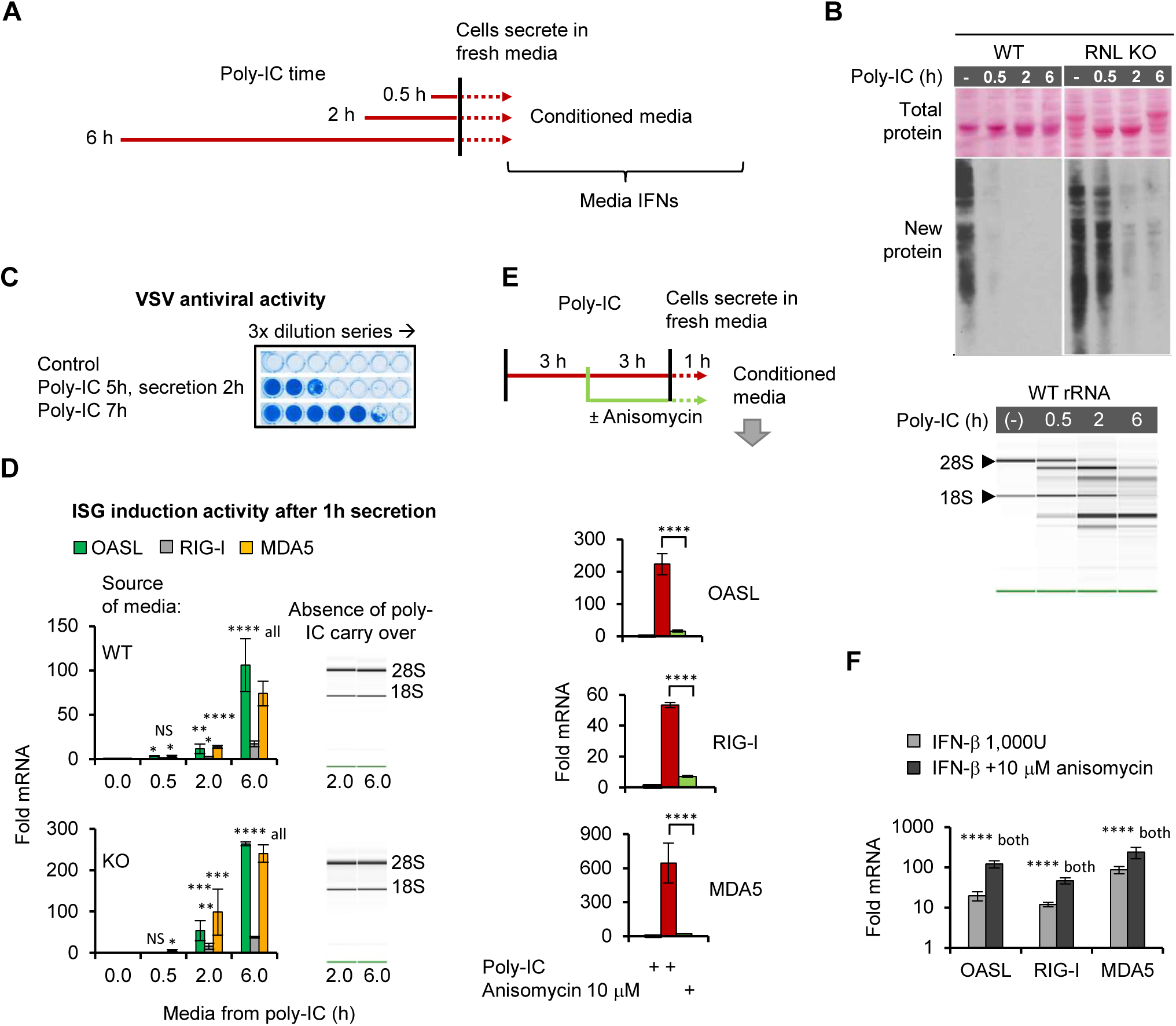
IFN synthesis after 2-5A-induced global translation shutoff. (**A**) Diagram of interferon secretion experiment. (**B**) Puromycin western blot (upper) and RNA nano-chip (lower) analysis of translation and 28S rRNA cleavage in poly-IC-treated A549 cells. Images are representative of four independent experiments. (**C**) Antiviral activity of conditioned media from poly-IC-treated A549 cells (upper). Condensed results from three additional replicates are shown on figure S8. (**D**) qRT-PCR analysis of ISG expression in WT and RNL-KO A549 cells treated with conditioned media. from poly-IC-treated A549 cells (lower). Data are means ± S.E. pooled from 3 biological replicates. RNA nano-chip (inset) of intact rRNA is representative of all experiments. (**E**) qRT-PCR analysis of ISG expression in A549 cells treated with anisomycin after translational arrest by poly-IC, but before the transcriptional IFN response. Data are means ± S.E. pooled from 3 biological replicates. (**F**) Effect of anisomycin treatment on transcriptional IFN signaling.

To test whether IFN arises from actively ongoing translation rather than from other potential mechanisms (e.g. delayed secretion of pre-translated IFN stores), we used pulse-treatment with a translation inhibitor, anisomycin (Fig. 5E). In this setting, cells were first treated with poly-IC for three hours, which stopped protein synthesis but did not yet activate a strong transcriptional IFN response. Next, anisomycin was added to arrest all protein synthesis and the cells were kept for three additional hours. Control cells were kept for the same duration without anisomycin. During the last hour, media was changed to remove poly-IC, but anisomycin treatment was continued to keep the cells translationally arrested. When IFN activity in the media was assayed, we found that anisomycin treatment after the 2-5A-induced global translational inhibition (fig. S8C), but before the IFN response, blocked IFN production (Fig. 5E). A control experiment showed that anisomycin was compatible with IFN sensing by naïve cells (Fig. 5F). Of note, anisomycin had a mild stimulatory effect on ISG mRNAs due to an unknown mechanism; this effect acted in the opposite direction from blocking IFNs and thus did not affect the suitability of anisomycin as a control in our tests. Together, our experiments indicated that IFNs are indeed translated when the bulk of protein synthesis remains silenced by 2-5A.

A549 cells treated with poly-IC express both type I and type III interferons (Fig. 6A). To determine which of these IFNs escape RNase L in our experiments, we employed hamster CHO reporter cell lines developed previously for specific detection of human IFNs of type I and type III. Hamster cells do not respond to human IFNs, however the reporter cells are rendered sensitive via expression of chimeric type I and type III human IFN receptors fused to a potent STAT1 docking domain^41,42^. The reporter cells analysis, based on readout of phospho-STAT, indicates the presence of type I and type III IFNs, exhibiting strongest p-STAT response to IFNs-λ (Fig. 6B and fig. S10).

**Fig. 6.**
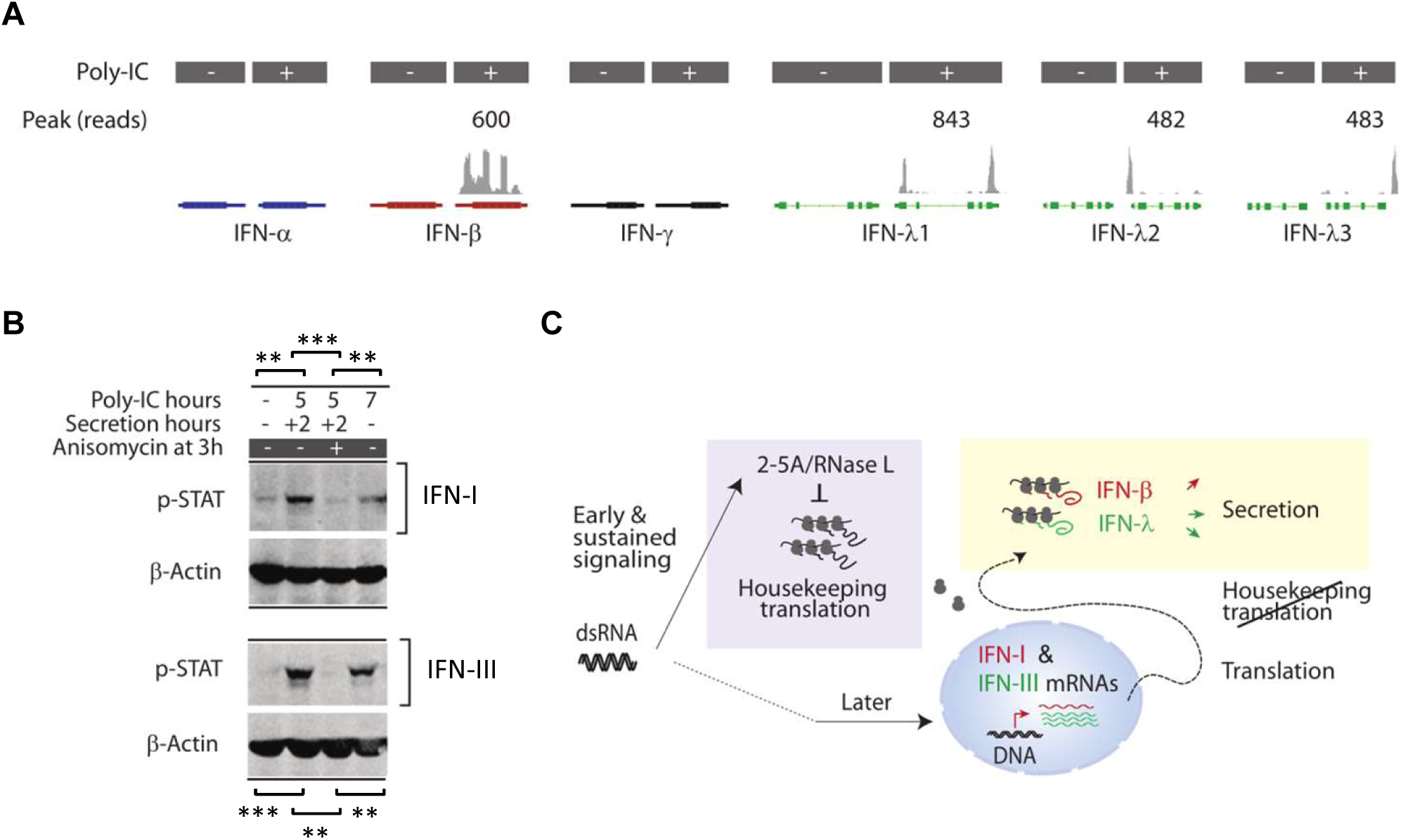
Type-I and type-III IFNs escape RNase L. (**A**) Poly-A^+^ RNA-seq profiles analysis of IFN mRNA expression in A549 cells treated with poly-IC (1 μg/ml, 9 hours). Data were mapped to hg19 assembly and plotted. Of note, our RNA-seq found that actual IFN-λ genes span slightly beyond their annotated coordinates in the reference genome hg19. This is still uncorrected in hg38. (**B**) Western blot analysis of pSTAT1 levels in CHO reporter cells for type I IFN (upper) or type III IFN (lower). Reporter cells were treated with conditioned media from A549 cells incubated with poly IC and anisomycin, as indicated. Blots are representative of 3 independent experiments. (**C**) Proposed role for 2-5A/RNase L in dsRNA sensing. 2-5A rapidly switches translation from basal proteins to prioritized IFN-β and IFN-λ synthesis and secretion.

## Discussion

We have developed a biosensor for 2-5A and determined that this second messenger is synthesized without a delay and mediates immediate dsRNA sensing. 2-5A activates RNase L, which suppresses protein synthesis^43^. This mechanism is potent and attenuates basal cell-wide translation by more than 1,000-fold (Fig. 4G, 5B and fig. S7B). Using this biosensor, we noted that 2-5A arrests host translational activity prior to induction of the interferon response. The translation-arrested cells still maintain efficient production and secretion of IFNs β and λ (Fig. 6C). Our work thus kinetically separates RNA cleavage and translational shutoff by RNase L from interferon-mediated cellular reprogramming, revealing an unanticipated order of signaling events where basal translation is shut down first and the IFN response develops second.

The action of RNase L resembles arrest of the initiation step, which can be accompanied by a characteristic collapse of polysomes^25^. A similar polysomal collapse is observed upon activation of integrated stress response (ISR) that employs serine/threonine kinases to phosphorylate and inactivate the translation initiation factor eIF2α^44^. In the ISR, the arrest of translational initiation can be bypassed by mRNAs encoding stress proteins, such as ATF4 and IBTKα due to the presence of 5’ uORFs that increase translation under conditions of limiting initiation^45^. Although it remains unclear how IFNs bypass RNase L and whether they may use a related route^46^, by evading 2-5A/RNase L translational arrest cells may ensure that infection does not prevent the production of IFNs, a central task of the innate immune system. In mice RNase L amplifies IFN protein synthesis^36^, indicating that cellular resources released by RNase L upon translational shutoff (translation can consume as much as 75% of a cell’s energy balance^47^) may become reallocated for enhanced production of IFNs.

The observations of subcellular 2-5A dynamics suggest a possible explanation for the bipartite organization of the OAS-RNase L system. The effector (RNase L) and the dsRNA-sensing moiety (the OASs) in the 2-5A system are separated. This arrangement contrasts with the single-protein structure of another dsRNA sensor, PKR, which encodes the dsRNA-binding domain and the effector kinase domain in the same polypeptide. Our data indicated that the bipartite arrangement of OASs/RNase L may sense dsRNA at a distance from the site of RNase L action. The range of OAS-RNase L communication depends on efficiency of 2-5A diffusion, which occurs with rates sufficient for 2-5A equilibration between the nucleus and the cytosol faster than the rate of 2-5A production. Therefore, RNase L is poised to respond to 2-5A from cytoplasmic and nuclear OASs, suggesting that detection of dsRNA in both compartments is a likely biologic function of cytosolic RNase L.

A number of clinically important translation inhibitors, such as rapalogs and a new generation of anticancer drugs based on INK128, work by reprogramming protein synthesis through inhibition of mTOR^48^. Our work for the first time describes RNase L not as a general RNA decay machine, but as a translation-reprogramming receptor. Normally, RNase L is activated as a part of the innate immune system. However, RNase L activation by small molecules could be explored for developing adjuvants and anticancer therapeutics with some of the beneficial effects of mTOR blockers, and with an added advantage of maintaining the protein synthesis activity of the innate immune system. The search for such on demand activators can be facilitated by biochemical, cell-based as well as in *vivo* applications of the 2-5A biosensor we describe here.

## Materials and Methods

### Tissue culture

Cells were grown using ATCC (American Type Culture Collection) or provider recommended conditions in MEM media + 10% FBS (HeLa) or RPMI media + 10% FBS (A549), or DMEM media + 10% FBS (293T). All media were purchased from Gibco, Life Technologies. HeLa and 293T were a gift from Yibin Kang (Princeton University, Princeton). WT, RNase L KO, and OAS KO A549 were generated by the laboratory of Susan Weiss (University of Pennsylvania, Philadelphia). Luminescence assays in live cells were carried out in a plate reader or in 12-well plates at 37 °C.

### Reporter protein design and preparation

Two halves of firefly luciferase encoding amino acids 1-416 (NFluc) and 417-550 (CFluc) were fused to the N- and C-termini of ANK domain of human RNase L (residues 21-325). For cell work, an N-terminal FLAG tag was added. PKKKRKVE and LQLPPLERLTLD were the NLS and NES sequences cloned immediately after the N-terminal FLAG tag. The reporter construct was synthesized by total DNA synthesis and cloned into pUC57 plasmid by GeneWiz. The construct was subcloned from pUC57 into pGEX-6P vector (GE Healthcare Life Sciences), which has N-terminal GST to aid purification.

GST-tagged reporter protein was expressed in *E.coli* BL21 (DE3) CodonPlus RIPL (Agilent technologies). Cells were lysed on Emulsiflex C3 (Avestin) in buffer containing 20 mM HEPES (pH 7.4), 300 mM NaCl, 5 mM MgCl_2_, 0.1 mM EDTA, 10% (v/v) glycerol, 5 mM DTT and 1% Triton X-100. Crude lysates were centrifuged at 35,000 g for 30 minutes. Clarified lysates were affinity purified using glutathione sepharose (GE Healthcare Life Sciences) and the GST tag was removed with Prescission Protease (GE Healthcare Life Sciences). Protein concentration was determined by band densitometry of coomassie-stained SDS PAGE gels using bovine serum albumin as a standard.

### Luminescence measurements using recombinant reporter

The luminescence reactions contained 20 mM HEPES (pH 7.4), 100 mM NaCl, 5 mM MgCl_2_, 5% (v/v) glycerol, 1 mM DTT, 3 mM ATP, 250 μΜ coenzyme A hydrate (Sigma Aldrich). After adding reporter protein and 2-5A (ppp-A_2,5_A_2,5_A) at concentrations specified on the figures, D-luciferin, potassium salt (Gold Biotechnology) was added at a final concentration of 400 μM. The reactions were allowed to stabilize for two minutes and luminescence readings were done using a Berthold Technologies microplate reader. Acquisition time for the luminescence was 10 seconds.

### 2-5A extraction from human cells and RNase L cleavage measurements

HeLa cells in 10 cm plates at 80% confluency were treated with 1 μg/mL poly-IC for 6 hours and lysed with 1 ml RIPA buffer (Thermo Fisher Scientific) supplemented with 0.5 mM PMSF protease inhibitor. The lysate was spun down at 12,000 g for 10 minutes at 4°C. The supernatant was passed through a 3 kDa centrifugal filter to collect the fraction of small nucleic acids. Equivalent amounts of small RNA from mock and poly-IC treated samples were used in the luminescence assays. 2-5A activity was tested using ^32^p-5’-radiolabeled RNA substrate and recombinant human RNase L as described previously^22^.

### Western blots

The V6 and V6-Y312A reporters were subcloned into pCDNA4.TO using primers to add an N-terminal FLAG tag. The constructs (2 μg of plasmid DNA) were transfected with 6 μl Lipofectamine 2000 (Thermofisher) into HeLa cells at 80% confluency. Twenty four hours later the cells were lysed in sample buffer (NuPage), separated on 10% BisTris PAGE (NuPAGE), and transferred to PVDF membranes (Life Technologies). Membranes were blocked in 5% nonfat dry milk in TBST for 30 min and probed with 1:2000 mouse anti-FLAG M2 (Sigma) or 1:5000 mouse anti-human GAPDH (Sigma) primary antibodies, at 4 °C overnight. The membranes were then washed with TBST and incubated with horseradish peroxidase-conjugated anti-mouse secondary antibodies (1:10,000 Jackson ImmunoResearch) for 30 minutes. The membranes were washed again and detected with ECL Western Blotting Detection Reagents (GE Healthcare Life Sciences) on an X-ray film. Western blots were quantified using local background algorithm in GelQuant.NET (http://biochemlabsolutions.com) and either ponceau/coomassie or ACTB as loading controls.

### Stable human cell lines expressing FLAG-V6 and FLAG V6-Y312A

For generating lentivirus, 293T cells were seeded into 6 well plates to achieve 50% confluency at 24 hours. Cells were transfected using FuGene with 1.5 μg pLEX.MCS (vector plasmid containing FLAG tagged WT or Y312A reporter), 1.33 μg pCMVdR8.91 (Gag-Pol packaging plasmid) and 0.17 μg pMD2.G (envelope plasmid). Lentivirus-containing medium was collected after 48 hours. Following collection, the medium was passed through 0.45 μm filter. Polybrene 5 μg/mL (f/c) and HEPES (pH 7.5; 100 mM f/c) were then added.

HeLa cells at 40% confluence were infected with 600 μl of lentivirus-containing media in 10 cm dishes. The media was changed after 24 hours post-infection, and puromycin (1.5 μg/mL f/c) was added 3 days post-infection. Monoclonal cells were picked by limiting dilution. For HeLa cells, single cell clones were screened based on high fold-changes in reporter activity during poly-IC treatment. A549 cells at 90% confluency were plated in 24 wells and transduced with 200 μl lentivirus. 48 hours post transduction, media containing 2 μg/mL puromycin was added. After 72 hours of selection, surviving cells were plated by limiting dilution to pick single clones. For A549 cells, single cell clones were screened by western blotting with anti FLAG M2 (Agilent).

### Live cell 2-5A measurements with transient transfection of reporter constructs

Cells were seeded on TC-treated, clear flat bottom white 96-well plates at 80% confluency. After the cells adhered to the plates (in approx 6 hours), 0.2 μg V6 or V6-Y312A (with or without localization tags) plasmids were transfected using 0.5 μL Lipofectamine 2000. 24 hours after transfection, the cells were pre-treated with 100 μΜ f/c D-luciferin ethyl ester (1% DMSO) (Marker Gene Technologies). After one hour of substrate pre-treatment, a poly-IC/lipofectamine complex (1 μg/mL Poly-IC + 0.5 μL of lipofectamine 2000 in a final volume of 100 μL) was added to the cells, supplemented with 100 μΜ D-luciferin ethyl ester substrate and 20 mM (f/c) HEPES (pH 7.5). Luminescence measurements were taken by Berthold Technologies plate reader in repeat-mode. Measurements were done every 15 minutes for three hours. Luminescence acquisition time was 1 minute per well.

### Immunofluorescence microscopy

HeLa cells were seeded in 8-chamber wells (Nunc™ Lab-Tek™ Chambered Coverglass) at 60% confluency. Cells were fixed with 4% paraformaldehyde for 15 minutes at room temperature after 24 hours post transfection of reporters (0.2 μg reporter plasmid and 1 μL Lipofectamine 2000). Cells were permeabilized with 0.1% Triton for 20 minutes at room temperature then blocked in 20% goat serum for 1 hour at 4 °C. The specimens were then incubated with 1:400 mouse anti-FLAG M2 (Sigma) primary antibodies for 2 hours at 4 °C and then rinsed 3x with excess PBS and incubated with AlexaFluor 488 goat anti-mouse (1:400). Stained cells were imaged using confocal microscope at 60x magnification.

### Live cell 2-5A measurements in stable cell lines expressing V6 or V6-Y312A reporters

A549 or HeLa cells stably expressing V6 or V6-Y312A control reporters were seeded at a density of 1⋅10^4^/well on TC-treated, clear flat bottom white 96-well plates. Before transfection with poly-IC, cells were pre-treated with 100 μM D-luciferin ethyl ester (1% DMSO) (f/c) (Marker Gene Technologies). After one hour of substrate pre-treatment, a poly-IC/lipofectamine complex (1 μg/mL Poly-IC + 0.5 μL Lipofectamine 2000 in a final volume of 100 μL) was added to the cells along with a second addition of 100 μΜ D-luciferin and HEPES (pH 7.5; 20 mM f/c). Dilutions of poly-IC were made by diluting the poly-IC/lipofectamine complexes. Luminescence measurements were taken in repeat-mode using plate reader at time intervals shown on the figures. After three hours, fresh D-luciferin ethyl ester (100 μΜ) was re-supplied at every hour to ensure excess reporter substrate. Luminescence acquisition time was one minute per well. For IFN-β pre-treatment, a dose of 1000 U/mL (f/c) was used 24 hours before poly-IC treatment.

### qRT-PCR analysis

Cells were harvested in 350 μL RLT buffer (Qiagen) and RNA was purified according to the RNeasy protocol (Qiagen). cDNA was prepared using oligo-dT and a High Capacity RNA to cDNA kit (Applied Biosystems). qPCR was performed using the Power SYBR green PCR mix in a 96 well format on StepOnePlus qPCR instrument (Life Technologies). qPCR primers used in this work were from (IDT). Primer sequences are listed in Table S1.

### Ribopuromycilation and ^35^S labeling to monitor nascent translation

To generate puromycin-tagged nascent peptides, human cells were treated with 0.1-1 μg/mL of poly-IC for times specified on the figures, after which the growth media was supplemented with 10 μg/mL puromycin (Invitrogen). Puromycin pulse lasted for 5 min. Cells were trypsinized and harvested in NuPAGE LDS sample buffer for western blot analyses. Proteins were separated by 10% BisTris PAGE (NuPAGE), and transferred on PVDF membranes (Life Technologies). The membrane was stained with Ponceau to normalize for sample loading, then washed and blocked with 5% non-fat dry milk in TBST buffer. The membranes were probed with 1:1000 mouse anti-puromycin antibody (EMD Millipore) that binds to *de novo* synthesized proteins, followed by horseradish peroxidase-conjugated goat anti-mouse secondary antibody (1:10,000, Jackson ImmunoResearch).

Metabolic labeling with ^35^S was conducted using the following procedure. After cell treatment with poly-IC, media was changed to methionine-free RPMI + 10% FBS supplemented with 11 μCi EasyTag™ EXPRESS35S Protein Labeling Mix (Perkin Elmer). Cellular proteins were resolved by 10% BisTris PAGE (NuPAGE) and analyzed by phosphorimaging.

### Secreted IFN detection by qPCR in conditioned media

A549 WT and RNase L KO cells in 12-well plates were transfected with 1 μg/mL poly-IC and lipofectamine 2000 for 0.5, 2 or 6 hours. At the end of the time course, cells were washed four times with 1 mL of growth medium (RPMI+10% FBS) to remove residual poly-IC. After the washes, fresh 1 mL medium was added and the cells were kept in fresh media for one hour to allow for protein secretion.

Treatments of naive cells with conditioned media were done in a separate 12-well plate and using cells seeded 1 day before the analysis. Naïve cell media was replaced with 1 mL of the conditioned medium from above. Cells were grown in the conditioned media for 16 hours and harvested in 300 μL RLT (Qiagen). For pulse-chase experiments in Fig. 5C, 10 μΜ anisomycin was added 3 hours after poly-IC addition. Anisomycin level was additionally maintained during the last hour allocated for IFN secretion into fresh media. To exclude a possible inhibitory effects of anisomycin on IFN response (Fig. 5D), WT cells were treated for 16 hours with 1000 U/mL recombinant IFN-β in plan or conditioned media.

### RNA-seq

Poly-A^+^ RNA sequencing was conducted and processed as described previously in our work^18,46^. The datasets (Fig. 5C) were deposited to GEO database under accession number GSE120355.

### Reporter assay to detect type-I and type-III IFNs

Previously developed reporter cell lines, which are selectively sensitive to either human type I^41^ or type III^42^ IFNs, were used to detect the presence of IFNs in the media. Briefly, 4×10^6^ cells were treated with each sample and incubated at 37 °C for 20 min. The cells were then washed with PBS, lysed and analyzed by immuno-blotting with antibodies specific for phosphorylated STAT1 (pSTAT1; BD clone 14/P-STAT1 (RUO)).

### Media antiviral activity assay

Equal numbers of human retinal pigment epithelial ARPE-19 cells were plated in all wells of a 96 well plate on day 1. On day 2, the cells were then treated with 3-fold serial dilutions of samples, starting from the first wells on the left to the last wells on the right. The cells were then incubated for 24 hours to allow induction of IFN response and then challenged with vesicular stomatitis virus, keeping the virus concentration constant in all wells (1⋅10^3^ PFU/well). The virus-treated cells are then incubated for 48 hours and the cells not killed by the virus are visualized by staining with crystal violet.

### Diffusion calculations

First, we estimate diffusive relaxation rate in a simplistic cell without the nucleus. This will define how fast a non-uniform concentration of 2-5A should relax in a sphere the size of a cell. The slowest mode to relax will be the spherically symmetric one with a single peak at the cell center, and zero gradient at the cell membrane (reflecting boundary conditions). The diffusion equation with no sources or sinks is: ∂c/∂t = D⋅ ⊽^2^c, which in the case of spherical symmetry simplifies to ∂c/∂t = D⋅(1/r^2^)⋅∂/∂r(r^2^⋅∂c/∂r). This simplifies further if we make the substitution c = u/r, and use ∂c/∂r = (1/r)⋅∂u/∂r - (1/r^2^)⋅u and ∂c/∂t = (1/r)⋅∂u/∂t, yielding (1/r)⋅∂u/∂t = D⋅[(1/r^2^)⋅∂/∂r(r⋅∂u/∂r - u)], and finally ∂u/∂t = D⋅∂^2^u/∂r^2^. We can expand u in eigenfunctions of the right hand side, which are just sines and cosines, and the general solution for u(r,t) will be of the form: u(r,t) = Σu_k_⋅sin(k⋅r)⋅exp(-λ_k_⋅t) plus similar terms for cosines. Rather than solving for complete generality, we note that we want the slowest decaying mode for which c is finite at r=0 (eliminating cosines), and has a no-flux boundary condition at the sphere’s radius R. No flux at R implies ∂c/∂r = 0, which implies that ∂u/∂r⋅(1/u) = 1/r at r = R. Therefore, k⋅cos(k⋅R)/sin(k⋅R) = 1/R, i.e. tan(k⋅R) = k⋅R, and we need to solve this equation to obtain k. The lowest-k solution (slowest mode of diffusion) is k⋅R = 4.49. After defining k* = 4.49/R, we find the corresponding relaxation rate of this mode from the diffusion equation: (-λ_k_*)⋅u = -D⋅k*^2^⋅u, so that λ_k_* = D⋅k*^2^.

Up to this point, we have neglected contributions to the solution for u(r,t) that are zero when acted on by ⊽^2^, so we can add terms c_0_(t)⋅r or c_1_(t) to u(r,t), or equivalently terms c_0_(t) or c_1_(t)/r to c(r,t). Only the constant term satisfies continuity at the center of the sphere for c(r,t), as well as zero flux at the boundary, and this constant term does not decay in time. So the solution for c(r,t) at long times has the form: c(r,t) = c_0_ + (c*/r)⋅exp(-λ_k_*⋅t)⋅sin(k*⋅r). It approaches the constant c_0_ at very long times, with the leading spatially non-uniform term decaying at a rate: λ_k_* = D⋅k*^2^ = D⋅(4.49/R)^2^.

A HeLa cell radius (R) is ~ 20⋅10^−4^ cm. Diffusion coefficients of small macromolecules (< 10 kDa) in the HeLa cytosol and nucleus are ~ 0.2 of diffusion in free buffer^49^ and for 2-5A (~1-2 kDa) are extrapolated^49^ as ~ 0.2⋅5⋅10^−6^ cm^2^/s = 10^−6^ cm^2^/s. These parameters imply a rate constant λ^k^* = D⋅(4.49/R)^2^ = 10^−6^⋅(4.49/20⋅10^−4^)^2^ = 5 s^−1^, or a relaxation time of approximately 0.2 s.

To evaluate how the nuclear envelope changes the relaxation rate between cytoplasm and nucleoplasm, we need to examine the ratio of the sum of circumferences of the nuclear pores to the circumference of the nucleus^50^. This ratio is given by pore diameter times number of pores divided by nuclear diameter: 5.2 nm × 2000/20 microns ≈ 0.5. From the calculations of Zarnitsyn et al.^50^ for transport through a collection of pores contained within a disc-shaped region in a flat sheet, the above value of 0.5 corresponds to transport at ~ 25% of the free diffusion limit. Therefore, in the presence of the nuclear envelope, we estimate that the net relaxation rate will be ~ 0.25⋅λ_k*_ ~ 1 s^−1^, corresponding to a relaxation time of 2-5A between nucleus and cytoplasm of 1s.

## Supporting information

## Acknowledgements

We are grateful to Prof. Bonnie Bassler (Princeton University) for help with instrumentation for this project, to Dr. Wei Wang for supervising the Icahn Genomics Institute RNA-seq facility, to Gary Laevsky, Director of Confocal Microscopy, and members of Korennykh laboratory for critically reading the manuscript. We thank Prof. Andrei Korostelev for important comments on the manuscript.

## Author contributions

A.C. developed and applied the reporter and analyzed transcriptional responses. S.R. characterized translational regulation and IFN regulation by RNase L. J.D. carried out the initial design of the ANK-split luciferase construct. K.D. analyzed translation regulation by RNase L. Y.L. and S.R.W. made A549 reporter cell lines. S.K. and R.R.S. performed antiviral activity tests. N.S.W. conducted diffusion analysis. A.C. and A.K. wrote the manuscript. A.K. supervised the work.

## Data and materials availability

All data needed to evaluate the conclusions in the paper are present in the paper or the Supplementary Materials.

