## Supplementary material for "Realtime 2-5A kinetics suggests interferons β and λ evade global arrest of translation by RNase L"

Submitted to BioRxiv

**This PDF file includes:**

Figs. S1 to S10
Table S1

### Supplementary Materials

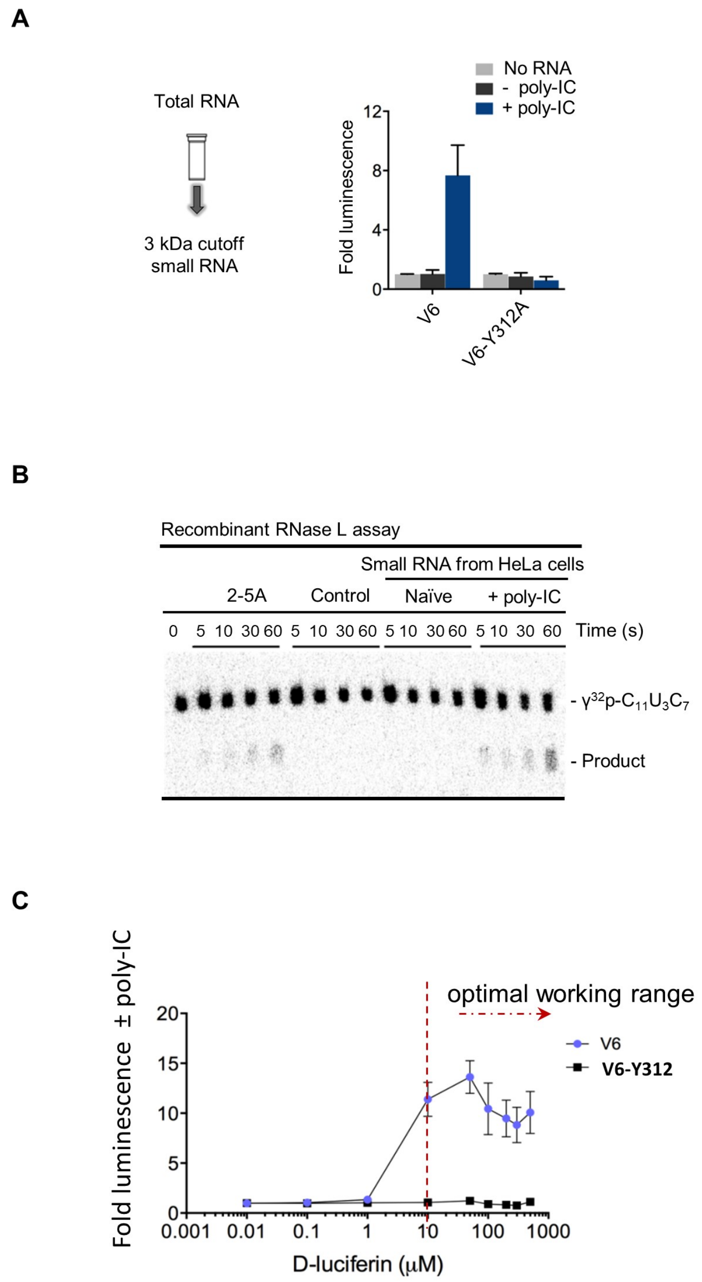


**Figure S1. Cellular 2-5A detection. A**) Detection of 2-5As extracted from cells treated with poly-IC for six hours. Equivalent $OD_{260}$ from each sample were used. Error bars represent S.E. from at least two independent experiments. **B**) RNA cleavage-based readout of cellular 2-5A using human RNase L (1 nM). **C**) Titration to define the working range of D-luciferin ethyl ester substrate. HeLa cells expressing V6 or V6-Y312A reporters were transfected with 1 μg/mL poly-IC for 6 h. Reporter luminescence was measured as described in Methods with using incremental doses of the luciferin substrate. Error bars show S.E. from three biological replicates.

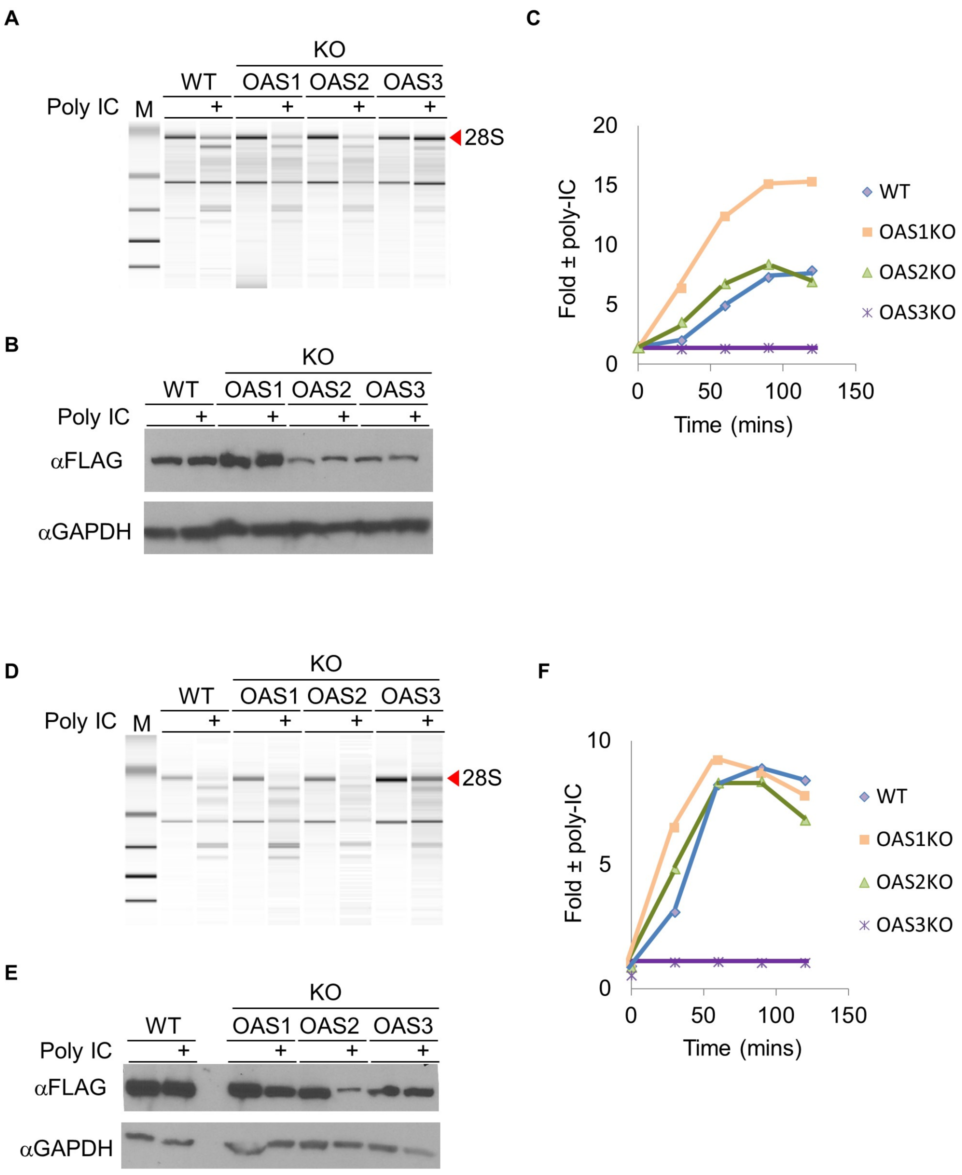


**Figure S2. Reporter luminescence response to poly-IC in OAS-KO A549 cells. A**) Bioanalyzer chip analysis of total RNA cleavage in FLAG-V6 reporter-expressing cells, control vs poly-IC-treated (1 µg/ml) for 2.5h. **B**) Western blot of FLAG-V6 protein in OAS KO cells. **C**). Reporter luminescence response in WT and cells with each OAS individually knocked out. **D-E**) An independent series of experiments as in A-C.

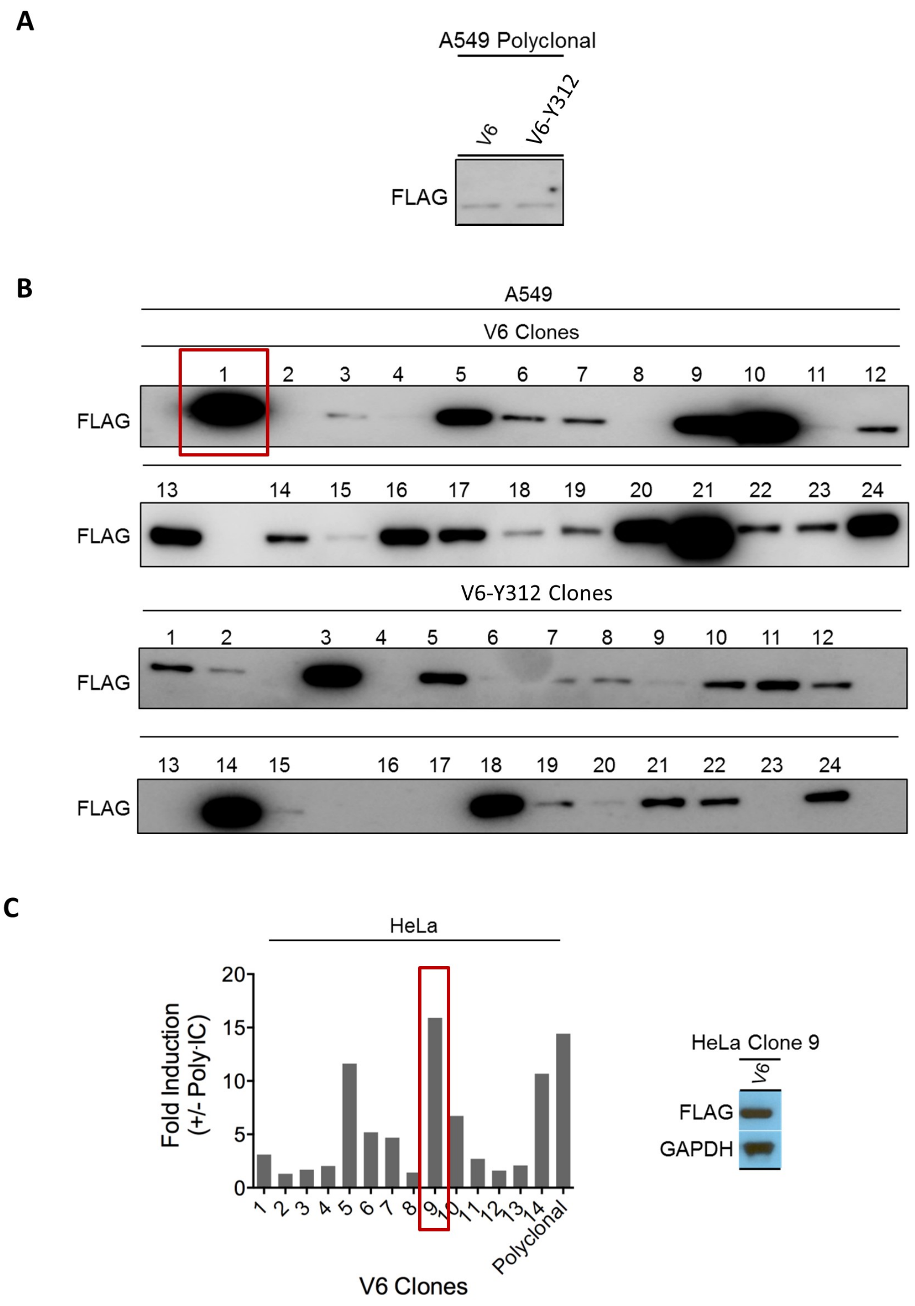


**Figure S3. Construction of A549 and HeLa cell lines stably expressing FLAG-V6. A)** Immunoblot analysis against FLAG tag in stably transduced polyclonal A549 cells. **B)** Immunoblot analysis against FLAG tag in monoclonal A549 cells. V6-Clone 1 was picked for all analyses in the main figures. **C)** Reporter activity induced by poly-IC treatment of single HeLa clones. Western blot analyses against FLAG tag in Clone 9. V6-Clone 9 was picked for all subsequent analysis.

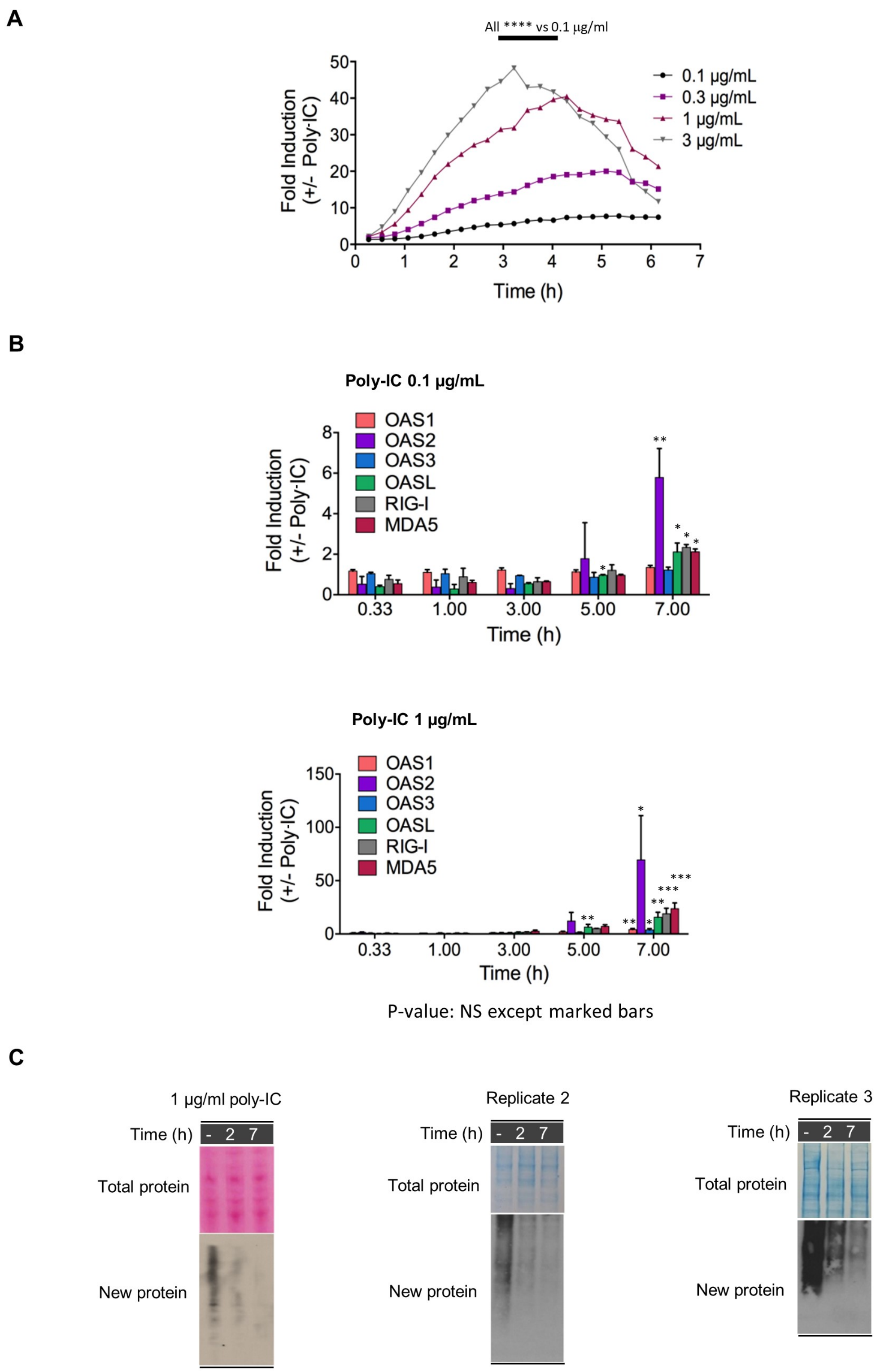


**Figure S4. 2-5A, transcription and translation profiling during poly-IC response in HeLa cells. A**) Reporter trace for 2-5A synthesis versus poly-IC concentration in HeLa cells that stably express V6 reporter. Luminescence assay was carried out as in Fig. 4A. P-values were calculated between 3h and 4h for three experimental replicates vs 0.1 µg/ml poly-IC. **B**) qPCR of select IFN-inducible genes in cells treated with 0.1 and 1 µg/mL poly-IC at multiple time pointes. Error bars show S.E. from either two or more biological replicates. **C**) Western blot analysis of nascent protein synthesis in naïve and poly-IC-treated HeLa cells.

A

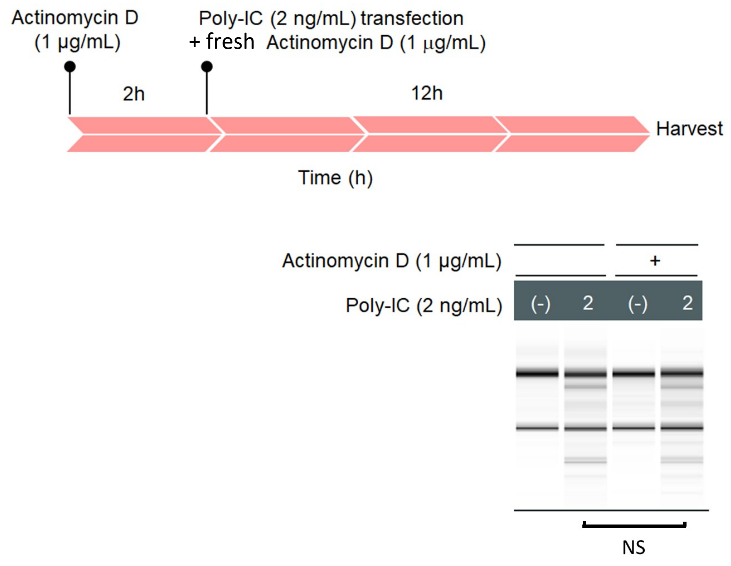


B

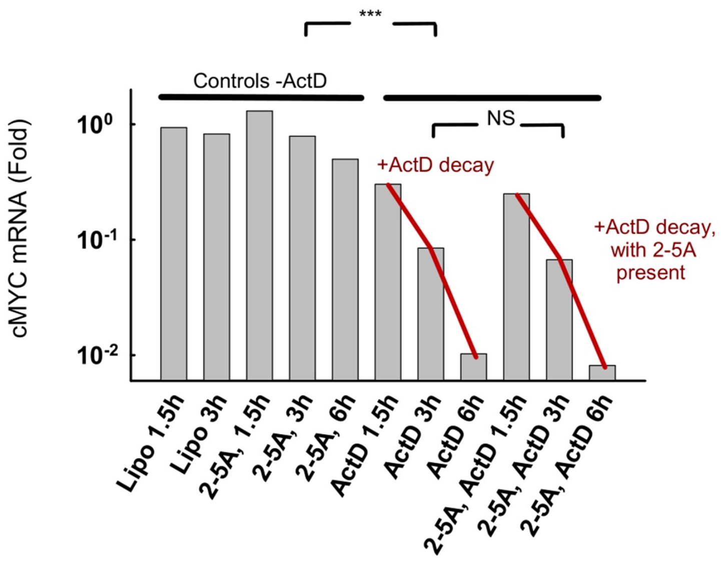


**Figure S5. Transcriptional response is not required for 2-5A synthesis. A)** Actinomycin D (ActD) pre-treatment for 12 hours does not preclude activation of the OASs/RNase L by poly-IC. P-value was computed using three experiments. **B**) ActD treatment works as determined by decay of cMYC mRNA via qPCR. Each bar represents an independent experiment. Statistical significance is shown for 5 input samples vs 6 experiments (***) and for three experiments "–A-25A" vs three experiments "+2-5A" (NS).

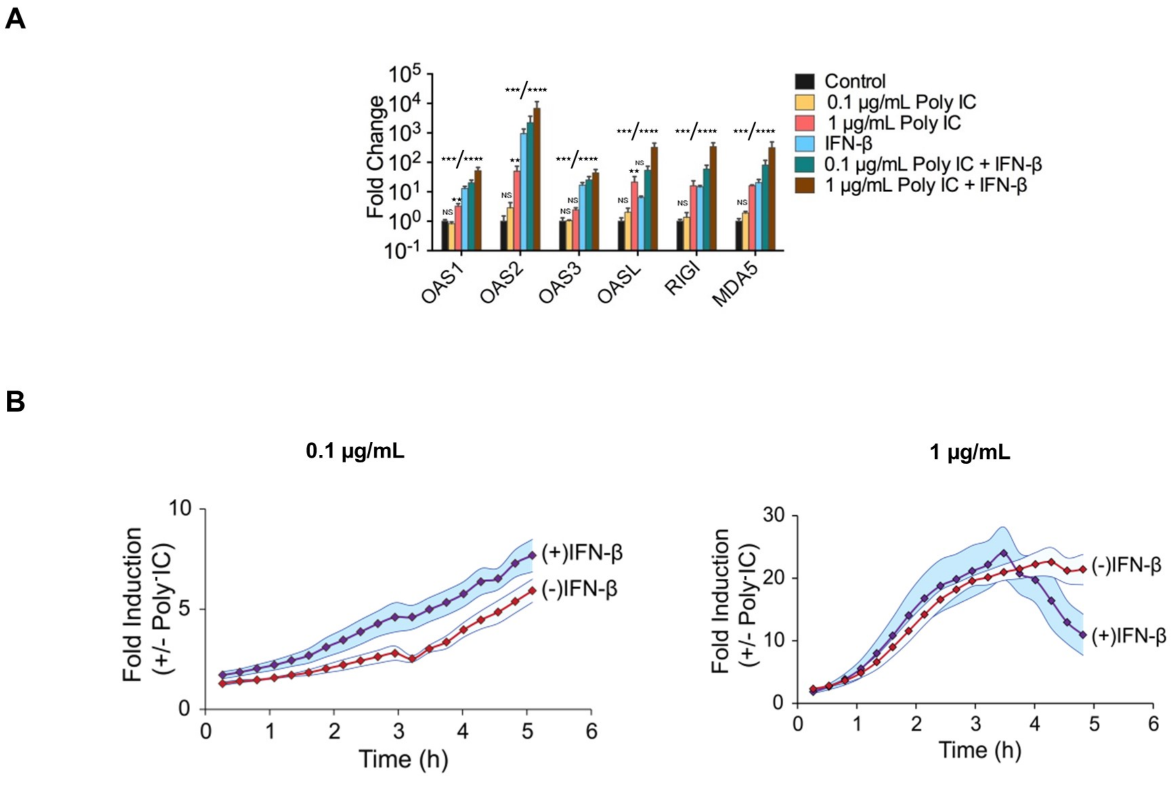


**Figure S6. 2-5A synthesis and transcriptional ISG induction mediated by poly-IC in IFN-β pre-treated HeLa cells. A**) Effect of 24 hours IFN-β pre-treatment (1000 U/mL) on transcriptional induction of ISGs in the presence of poly-IC. Poly-IC treatment was carried out for 5 hours. Errors are S.E. from three biological replicates. **B**) Poly-IC-induced reporter signal in HeLa cells stably expressing V6 with or without 24 hours of IFN-β pre-treatment (1000 U/mL). Shaded regions show S.E. from two biological replicates.

**A**

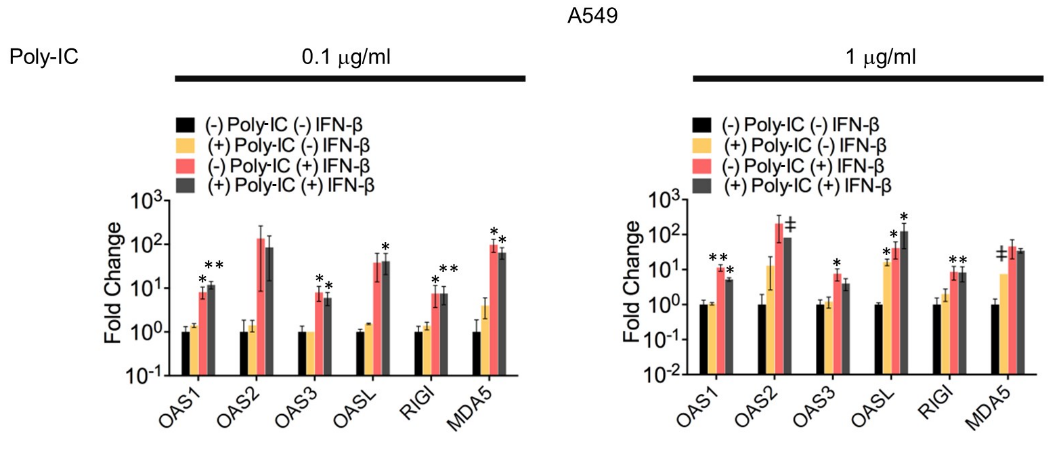


**B**

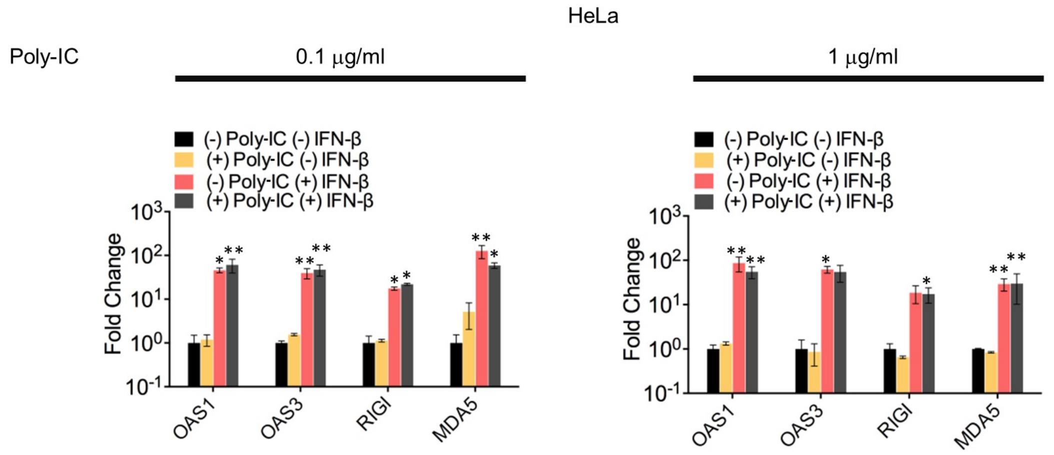


**Figure S7. Effect of IFN-β pre-treatment on transcriptional induction of ISGs in reporter-expressing stable A549 and HeLa cells.** Targeted qPCR of IFN inducible dsRNA sensors: the OAS1, OAS2, OAS3, OASL, and the helicases RIG-I and MDA5 in stable reporter cells: (A) A549; (B) HeLa. Reporter assays for these samples were carried out as in (Figs 3B and 4E), followed by RNA harvesting for qPCR analyses. Error bars represent S.E. from at least two biological replicates. Bar marked with ǂ is a single-replicate.

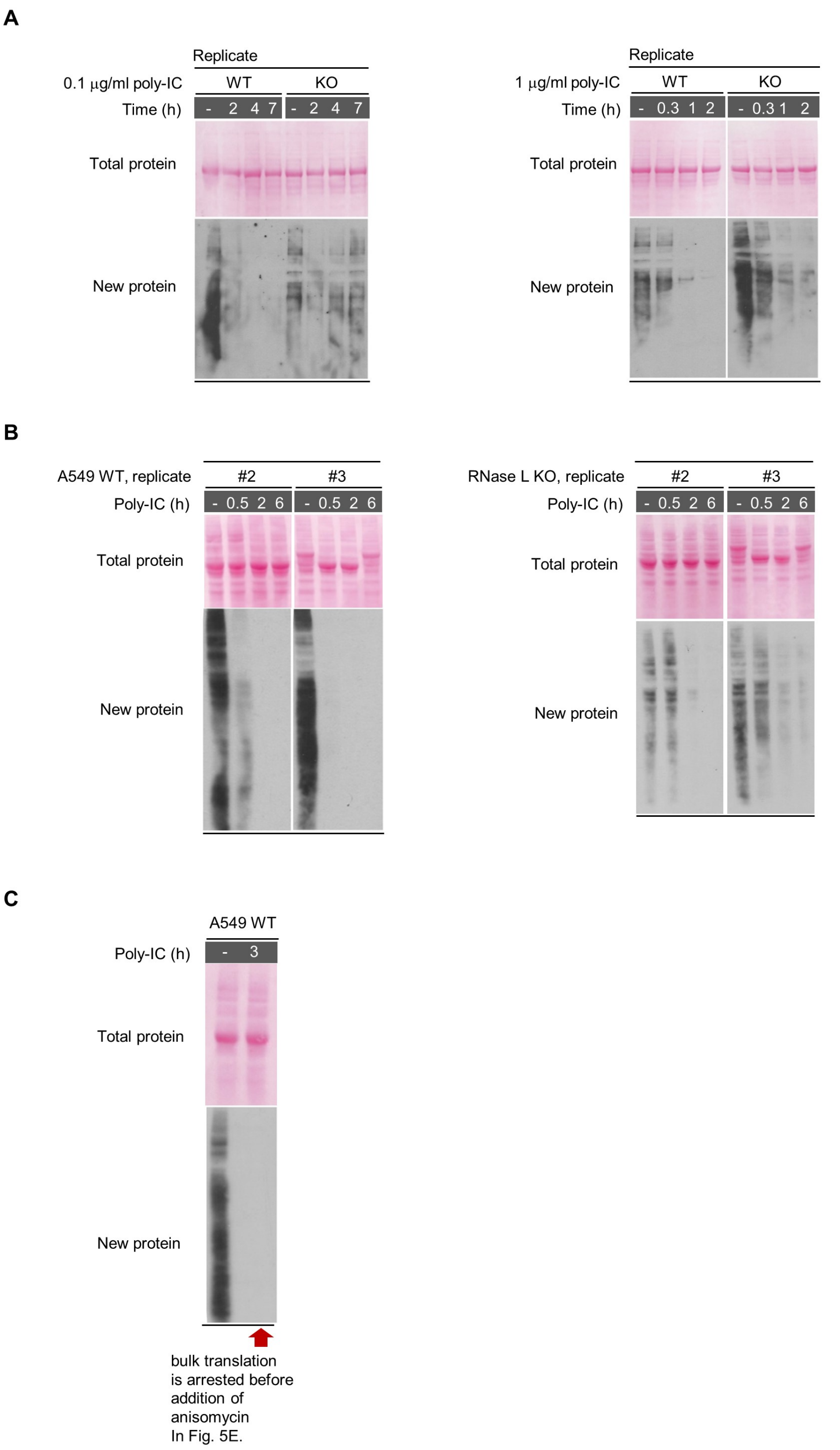


**Figure S8. Effect of poly-IC on global translation.** **A**) Biological replicates of the translational arrest in Fig. 4f. **B**) Biological replicates of the translational arrest in Fig. 5B. **C**) Western blot against puromycin-labeled nascent peptides to verify ongoing translational shutoff in Fig. 5E at three hours of poly-IC treatment (prior to adding anisomycin). Anisomycin dampens translation of IFNs that are synthesized after poly-IC-induced arrest.

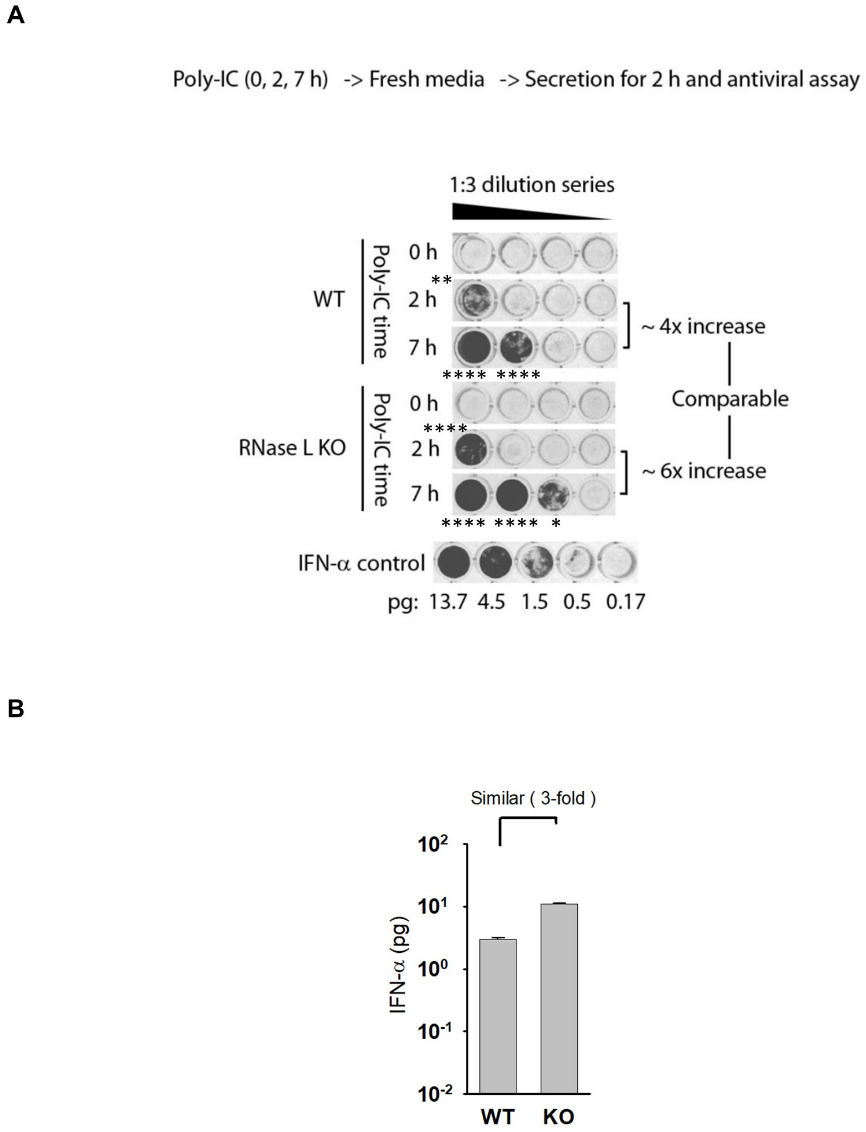


**Figure S9. Secreted media IFN measurements.** A) Conditioned media antiviral activity with VSV virus (Methods). Media from poly-IC-treated cells was collected and assayed for antiviral defense as in Figure 5C. These measurements show that WT and RNase L-KO A549 cells secrete comparable quantities of IFNs. Both cell types continue making IFNs after 2 hours of poly-IC treatment, where RNase L blocked all visible translation. Images and statistical quantification are representative of three (WT) and two (KO) biological replicates. B) Quantitation of the data in A signal using IFN-α calibration curve. Error bars show S.E.

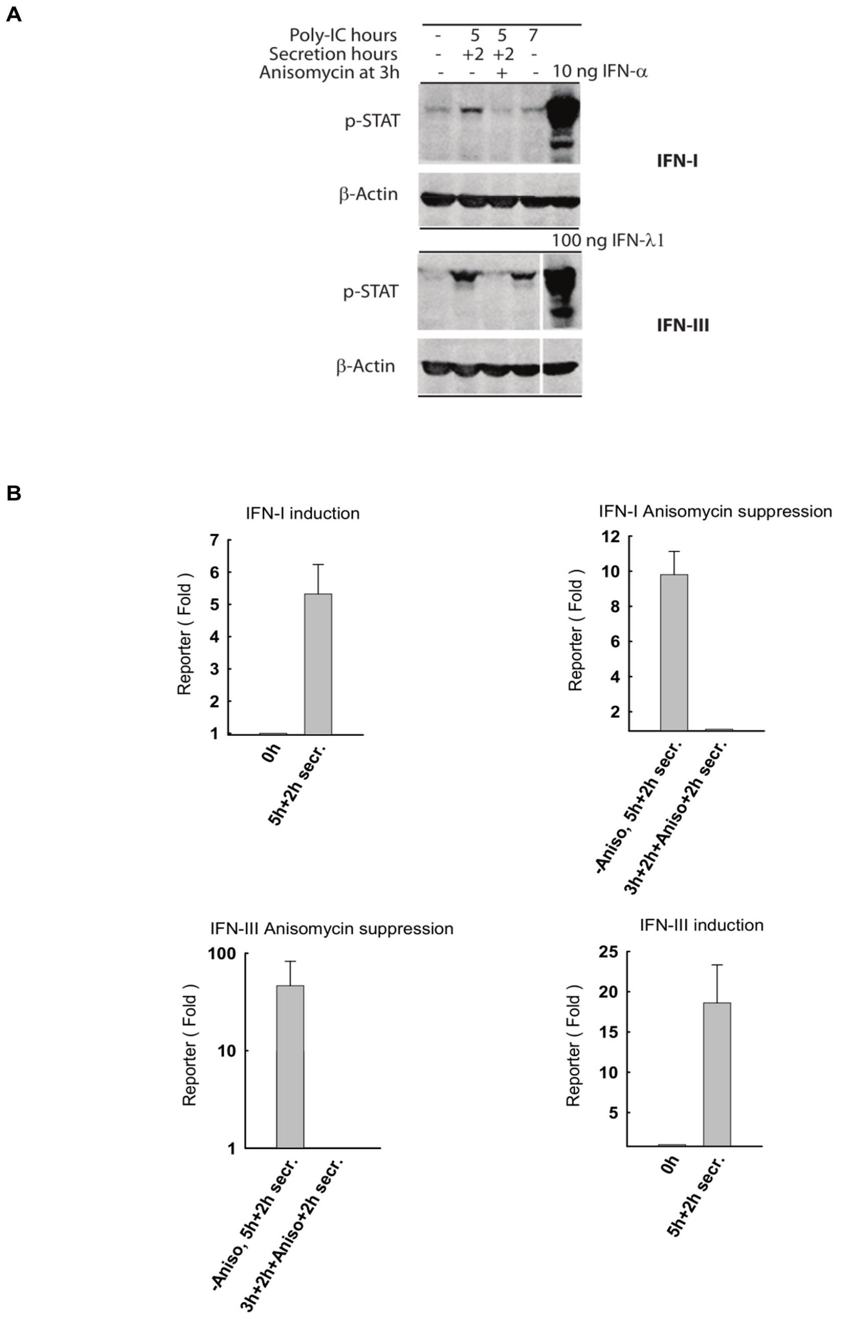


**Figure S10. Media IFN type analysis.** A) An independent measurement of type-I and type-III IFN activity in media with hamster reporter cells as in Fig. 6B. Assay was done in 96-well plate with 100 µl media volume per well. B) Quantitation of the gel. Error bars show S.E. from two independent series of experiments.

**Table S1.** qPCR Primers used

| Gene | Forward | Reverse |
|---|---|---|
| MDA5 | GCTTCTAGTTAG | CTTACACCTGATTCATTTCC |
| RIG-I | AGGAACTGGAGC | AGACTCTCTGTGTCCCTCAT |
| ACTB | CACTCTTCCAGC | GTACAGGTCTTTGCGGATG |
| OAS1 | Hs.PT.58.19958183 | |
| OAS2 | Hs.PT.58.24570700 | |
| OAS3 | Hs.PT.58.45631304. | |
| OASL | HS.PT.58.50426392 | |
